# Estimating heritability without environmental bias

**DOI:** 10.1101/218883

**Authors:** Alexander I. Young, Michael L. Frigge, Daniel F. Gudbjartsson, Gudmar Thorleifsson, Gyda Bjornsdottir, Patrick Sulem, Gisli Masson, Unnur Thorsteinsdottir, Kari Stefansson, Augustine Kong

## Abstract

Heritability measures the proportion of trait variation that is due to genetic inheritance. Measurement of heritability is of importance to the nature-versus-nurture debate. However, existing estimates of heritability could be biased by environmental effects. Here we introduce relatedness disequilibrium regression (RDR), a novel method for estimating heritability. RDR removes environmental bias by exploiting variation in relatedness due to random segregation. We use a sample of 54,888 Icelanders with both parents genotyped to estimate the heritability of 14 traits, including height (55.4%, S.E. 4.4%) and educational attainment (17.0%, S.E. 9.4%). Our results suggest that some other estimates of heritability could be inflated by environmental effects.

## Introduction

Heritability measures the proportion of trait variation in a population that is due to genetic inheritance. The relative importance of genetic inheritance (nature) versus environment (including nurture) for human traits has generated much controversy^1^. Historically, most estimates of heritability for human traits have come from twin studies^2,3^. More recently, GREML (genomic relatedness matrix restricted maximum likelihood) methods have been developed^4-8^. GREML methods propose to estimate heritability, or some fraction of it, by modelling the effects of genome-wide single nucleotide polymorphisms (SNPs). In order to reduce the influence of non-additive genetic effects and environmental effects, samples are pruned of close relatives before application of GREML methods.

Instead of modelling the effects of SNPs directly, heritability can be estimated by examining how phenotypic similarity changes with relatedness. Relatedness is measured by the fraction of the genome a pair shares in segments inherited from a common ancestor, called IBD (identical-by-descent) segments. (We note that what we call ‘relatedness’ here has sometimes been termed ‘realised relatedness’ to distinguish it from expected relatedness given a pedigree^9^). Sharing of an IBD segment implies sharing of all genetic variants in that segment, except for mutations that occurred since the last common ancestor of the segment. This implies that IBD based methods can capture nearly all of the heritability of a trait. In contrast, GREML methods can only capture the fraction of the heritability explained by genotyped SNPs^4^. Another advantage of IBD based methods over GREML methods is that they do not make assumptions about the distribution of SNP effect sizes. Violation of these assumptions has been shown to introduce bias to GREML estimates of heritability^4,10^.

An IBD based method, which we call the ‘Kinship’ method, examines how phenotypic similarity increases with relatedness for all pairs from a representative population sample^11^. When close relatives have more similar environments than distant relatives, the Kinship method will overestimate heritability, as it is unable to distinguish between similarity due to genetic effects and environmental effects. Another approach, which we call ‘Sib-Regression’, restricts the analysis to sibling pairs. Most of the variation in relatedness between siblings is due to random segregations in the parents, which are independent of environmental effects. Sib-Regression therefore avoids most sources of environmental bias. However, Sib-Regression requires hundreds of thousands of genotyped sibling pairs to obtain precise heritability estimates, whereas existing applications have used ∼20,000 sibling pairs or less^9,12^.

We introduce a novel method for estimating heritability, relatedness disequilibrium regression (RDR). RDR looks at how much more or less related a pair is than would be expected from the relatedness of the parents. This deviation we call ‘relatedness disequilibrium’. Relatedness disequilibrium is due to random segregation in the parents during meiosis, so is independent of most environmental effects. By using all pairs from a large sample with both parents genotyped, RDR can obtain precise estimates of heritability with negligible bias due to environment. We apply RDR to estimate heritability for 14 quantitative traits in Iceland.

## Results

### Defining heritability through random segregation

To define the heritability of a trait, we first distinguish direct genetic effects and indirect genetic effects: a direct genetic effect is the effect of genetic material in a body on that body, whereas an indirect genetic effect is the effect of genetic material in a body on another body (Supplementary Note)^13-15^. For example, if parenting affects the educational attainment of offspring, then there could be indirect genetic effects from parent to offspring, which we term ‘parental genetic nurturing effects’^15^. Any allele inherited by the phenotyped individual (proband) was also present in one of its parents, implying the allele can have both direct and parental genetic nurturing effects on the proband. However, parental genetic nurturing effects, and other indirect genetic effects, are environmental effects from the perspective of the individual whose trait is affected. The heritability of the trait is thus defined as the fraction of trait variation in the population that is explained by direct genetic effects alone. This is different from the fraction of trait variation explained by variation in proband genotype, which can include variation due to indirect genetic effects from the proband’s relatives.

To separate variation due to direct genetic effects (heritability) from variation explained by the environment, including indirect genetic effects from relatives, we use random segregation during meiosis. This approach is analogous to the transmission disequilibrium test (TDT) for a direct genetic effect of an allele on a phenotype^16-18^. Proband genotype is determined by the genotypes of the proband’s parents and random segregations. The TDT looks for an association between the phenotype and the variation in proband genotype caused by random segregations in the parents. This separates association due to direct genetic effects from association due to environment. Similarly, by using random segregation, phenotypic variation can be decomposed into variation due to direct genetic effects alone and other components. Assuming direct genetic effects are additive and there is no gene-by-environment interaction, the decomposition is (Supplementary Note):

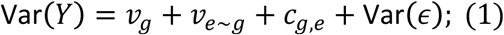

where *v*_*g*_ is the variance explained by direct genetic effects, and *h*^2^ = *v*_*g*_/Var(*Y*) is the heritability; *v*_*e∼g*_ is the variance of the part of the environmental component of the phenotype that is correlated with parental genotype, which includes the variance explained by (additive) parental genetic nurturing effects; *c*_*g,e*_ is the covariance between direct genetic effects and environmental effects; and Var(*∊*) is the variance of the component of the phenotype that is uncorrelated with both proband genotype and parental genotype.

### RDR covariance model

The variance decomposition (1) leads to a decomposition of the covariance matrix of a vector of phenotype observations, ***Y***. Under certain assumptions (Supplementary Note):

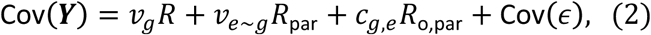

where [*R*]_*ij*_ is the relatedness of individual *i* and individual *j*, [*R*_par_]_*ij*_ is the relatedness of the parents of *i* and the parents of *j*; [*R*_o,par_]_*ij*_ is the relatedness of *i* and the parents of *j* and *j* and the parents of *i* (Methods). In general, Cov(*∊*) is unknown and can be similar to *R* due to population structure and/or environmental effects shared between relatives. To fit the RDR covariance model, we make the simplifying assumption that Cov(*∊*) = *σ*^2^I. Importantly, violation of the assumption that Cov(*∊*) = *σ*^2^I does not introduce bias to RDR estimates of heritability, as we outline below.

### RDR estimates heritability with negligible bias due to environment

The TDT separates direct genetic effects from environmental effects by using random segregation. RDR does something similar for variance components: by using random segregation, RDR separates the correlations between individuals’ phenotypes due to direct genetic effects from the correlations due to environmental effects. Just as the TDT conditions on parental genotype to remove bias due to association between genotype and environment, RDR conditions on parental relatedness to remove bias due to an increase in environmental similarity with relatedness. The expectation of offspring genotype given its parents’ genotypes is one half of the sum of the parents’ genotypes, and any variation around this expectation comes from random segregation. Similarly, the expectation of offspring relatedness, [*R*]_*ij*_, given parental relatedness, [*R*_par_]_*ij*_, is [*R*_par_]_*ij*_/2, and any variation around this expectation comes from random segregation (Figure 1, Supplementary Figure 1, and Supplementary Note). (Note that this relationship does not hold for pairs where one is the direct ancestor of the other, such as parent-offspring pairs.)

**Figure 1:**
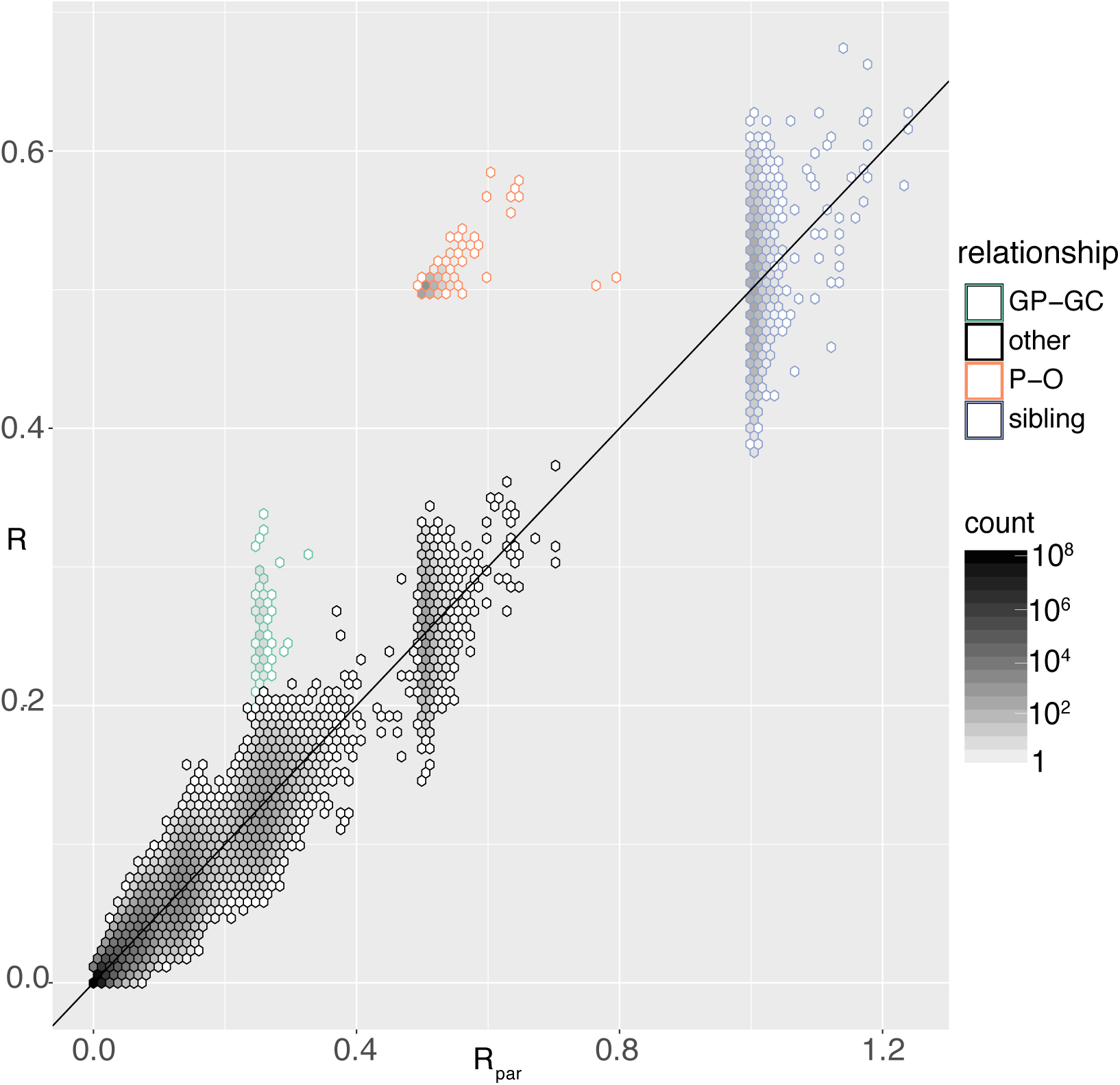
**Relatedness disequilibrium**: For all pairs of individuals *i,j* from 20,000 Icelanders with both parents genotyped, the relatedness of *i* and *j*, [*R*]_*ij*_, is compared to the relatedness of the parents of *i* and the parents of *j*, [*R*_par_]_*ij*_. The number of pairs in each hexagonal bin is indicated by shading. Relationships determined by the deCODE Genealogy database are indicated: GP-GC, grandparent-grandchild; P-O, parent-offspring; and sibling. The diagonal line indicates the expectation of [*R*]_*ij*_, which is [*R*_par_]_ij_/2, except for pairs where one is a direct ancestor of the other (Supplementary Note). Relatedness disequilibrium is the variation in [*R*]_*ij*_ around [*R*_par_]_*ij*_/2. Relatedness disequilibrium is due to independent, random segregations in the parents, except for pairs where one is the direct ancestor of the other.

By fitting *R* and *R*_par_ jointly, RDR uses the variation in [*R*]_*ij*_ around its expectation, [*R*_par_]_*ij*_/2, to estimate heritability. We call this variation relatedness disequilibrium. Relatedness disequilibrium is caused by random segregation in the parents, so is independent of sharing of all environmental effects apart from indirect genetic effects between *i* and *j*. This insight forms the basis of a mathematical proof that heritability estimates from RDR converge to the true heritability, when the sample excludes pairs that have indirect genetic effects on each other and excludes pairs where one is the direct ancestor of the other (Supplementary Note). If indirect genetic effects are restricted to close relatives, the bias is likely to be small for RDR because close relatives comprise only a small fraction of the pairs in a large population sample. The bias due to indirect genetic effects could be much larger for methods that rely on close relatives, such as Sib-Regression and twin studies.

Pairs where one is the direct ancestor of the other can introduce bias because they have an atypical relationship between [*R*]_*ij*_ and [*R*_par_]_*ij*_ (Figure 1). However, they will comprise a small fraction of the total pairs in a large population sample, even if multiple generations are genotyped. For our sample, around 30% also have a parent or grandparent in our sample, but parent-offspring and grandparent-grandchild pairs comprise only 0.0014% of all pairs. We performed simulations to detect bias due to inclusion of parent-offspring and grandparent-grandchild pairs in our sample (Methods). In our simulations, we could not detect any such bias in RDR heritability estimates (Supplementary Table 1), suggesting it is negligible for our sample. We therefore did not remove individuals from our sample that also have a parent or grandparent in our sample.

### Simulation of RDR heritability estimation

We simulated traits in our sample to demonstrate that RDR heritability estimates are approximately unbiased in situations where some other methods can be strongly biased. We simulated the genetic components of traits using the genotypes of 10,000 Icelandic individuals and their parents at 10,000 SNPs (Methods). The SNPs had a minimum minor allele frequency (MAF) of 0.5% and median MAF of 22.8%. We compared heritability estimates from RDR; the Kinship method; the Kinship method allowing for an effect of shared family environment, which we call the ‘Kinship F.E.’ method. (We determined whether pairs shared a family environment by whether they shared a mother according to the deCODE Genealogy Database.) We also compared to Sib-Regression but, to ensure we had enough sibling pairs, we simulated traits in the full sample of 54,888 individuals with both parents genotyped.

We first confirmed that heritability estimates for all the methods were approximately unbiased for traits determined by additive, direct genetic effects and random noise (‘additive’ trait, Table 1, Supplementary Tables 2 and 3).

**Table 1.** **Simulation results.** The mean heritability estimates, expressed as a % of the phenotypic variance, from four different methods (RDR, Kinship, Kinship F.E., Sib-Regression) for different simulated traits along with standard errors in brackets. The true (narrow-sense) heritability of each trait was 40%. We simulated 500 replicates of each trait based on actual Icelandic genetic data (Methods). Ten thousand SNPs with median minor allele frequency (MAF) 22.8% were given additive effects for all the traits other than the ‘rare SNPs’ trait, for which 2,200 SNPs with MAF between 0.1% and 1% (median 0.26%) were used. To the additive genetic component, only noise was added for the ‘additive’ trait and the ‘rare SNPs’ trait. For the epistatic trait, 10% of the phenotypic variance was due to pairwise interactions between SNPs. For the other traits, effects representing different sources of environmental confounding were added in addition to noise and the additive genetic component. For the ‘regional’ trait, each region of Iceland (sysla) was given an effect; for the ‘maternal environment’ trait, an environmental effect shared between those who share mothers was added; for the ‘genetic nurturing trait’, the genotypes of the parents were also given effects to simulate ‘parental genetic nurturing’ effects^15^. For the ‘regional’ trait, the Kinship and Kinship F.E. methods also included adjustment for 20 genetic principal components.

|  | RDR | Kinship | Kinship F.E. | Sib-Regression |
| --- | --- | --- | --- | --- |
| additive | 39.3 (0.62) | 40.4 (0.15) | 40.5 (0.18) | 41.2 (0.69) |
| genetic nurturing | 39.4 (0.49) | 92.7 (0.09) | 82.8 (0.14) | 40.4 (0.37) |
| maternal environment | 38.9 (0.58) | 76.3 (0.17) | 39.9 (0.18) | 41.1 (0.37) |
| regional | 38.3 (0.60) | 59.0 (0.17) | 58.3 (0.20) | 32.1 (0.63) |
| rare SNPs | 35.0 (0.64) | 39.5 (0.15) | 39.4 (0.19) | 39.7 (0.67) |
| epistatic | 41.3 (0.60) | 44.2 (0.16) | 43.3 (0.19) | 50.1 (0.63) |

We simulated a trait where individuals who shared a mother according to the deCODE genealogy database shared a random environmental effect. We found that the Kinship method greatly overestimated the heritability of this trait (‘maternal environment’ trait, Table 1). However, the Kinship F.E. estimates of heritability were approximately unbiased. Both Sib-Regression and RDR estimates were approximately unbiased.

The results for the ‘maternal environment’ trait show that modelling a family environment effect can remove bias from the Kinship method in certain circumstances. However, when indirect genetic effects from relatives are present, modelling the family environment is ineffective at removing bias. To show this, we simulated a trait determined by direct genetic effects, parental genetic nurturing effects, and random noise (‘genetic nurturing’ trait, Table 1). Let *δ* be the direct effect of a SNP, and let *η* be the genetic nurturing effect, which is the effect of parental genotype on offspring through the offspring’s environment. The total variance explained by parent and offspring genotype at the SNP is proportional to *δ*^2^ + 2*η*^2^ + 2*ηδ*, with *δ*^2^ proportional to the contribution to *v*_*g*_, 2*η*^2^ proportional to the contribution to *v*_*e∼g*_, and 2*ηδ* proportional to the contribution to *c*_*g,e*_. For the simulated trait, the genetic nurturing effect of each SNP was a fixed fraction of its direct effect, generating a substantial covariance term, *c*_*g,e*_. The variance components as a percentage of the phenotypic variance were: *v*_*g*_ = 40%, *v*_*e∼g*_ = 10%, and *c*_*g,e*_ ≈ 28%, bringing the total variance explained by parent and offspring genotype to ∼78%.

We found that the Kinship method greatly overestimated the heritability of the ‘genetic nurturing’ trait and that this bias was only slightly reduced by modelling a family environment effect. The reason for this is that parental genetic nurturing effects induce correlations between all pairs with non-zero parental relatedness, not just those that share a family environment. This leads to an increase in environmental similarity with relatedness across the relatedness spectrum. A similar increase in environmental similarity with relatedness would be induced by indirect genetic effects from other relatives, such as siblings.

Both RDR and Sib-Regression estimates of heritability were approximately unbiased for the ‘genetic nurturing’ trait. Furthermore, RDR estimates of *v*_*e∼g*_ and *c*_*g,e*_ were approximately unbiased estimates of the variance from parental genetic nurturing effects and the covariance between direct genetic effects and parental genetic nurturing effects (Supplementary Tables 2 and 3). In general, however, RDR estimates of *v*_*e∼g*_ and *c*_*g,e*_ will capture other sources of environmental variation in addition to parental genetic nurturing effects (Supplementary Note).

It has been recognized that population stratification causes bias in heritability estimates from GREML methods and the Kinship method^6,19,20^. We simulated a trait affected by population structure. For this trait, each region of Iceland had a different mean trait value (Methods). We found that the Kinship and Kinship F.E. estimates of heritability were upwardly biased even after adjusting for 20 genetic principal components (‘regional’ trait, Table 1). In contrast, RDR estimates were approximately unbiased.

In some cases, IBD based methods such as RDR will not capture the phenotypic variance explained by recent mutations, which are rare in the population. This is because, if a causal mutation arose after the last common ancestor of an IBD segment, sharing of that IBD segment is not informative of sharing of that mutation. To measure how well RDR captures variance from rare variants, we simulated a trait determined by additive, direct effects of SNPs with MAFs between 1% and 0.1%, with median MAF 0.26% (Methods). RDR captured ∼88% of the variance explained by the rare SNPs, less than the Kinship method, which captured ∼99% of the variance explained by the rare SNPs (‘rare SNPs’ trait, Table 1). This discrepancy is likely due to the fact that RDR gives less weight to information from close relatives than the Kinship method.

To investigate the sensitivity of RDR to the presence of genetic interactions, we simulated a trait with a narrow-sense heritability of 40% (40% of the phenotypic variance explained by additive effects of SNPs) and with an additional 10% of the phenotypic variance explained by pairwise interactions between SNPs (Methods). The mean RDR estimate of heritability was 41.3%, indicating a small upward bias due to genetic interactions. Sib-Regression is expected to estimate the sum of the narrow-sense heritability and the proportion of variance due to pairwise interactions, which the simulations confirm (‘epistatic’ trait, Table 1).

### RDR estimates of heritability for 14 human traits

We estimated the variance components of the RDR covariance model for 14 quantitative traits (Methods, Table 2, Supplementary Table 4, and Supplementary Figure 2). Let 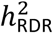 represent the RDR estimate of heritability. For height, 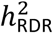 = 55.4% (SE = 4.4%). For educational attainment, 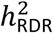 = 17.0% (S.E. = 9.4%).

**Fig. 2.**
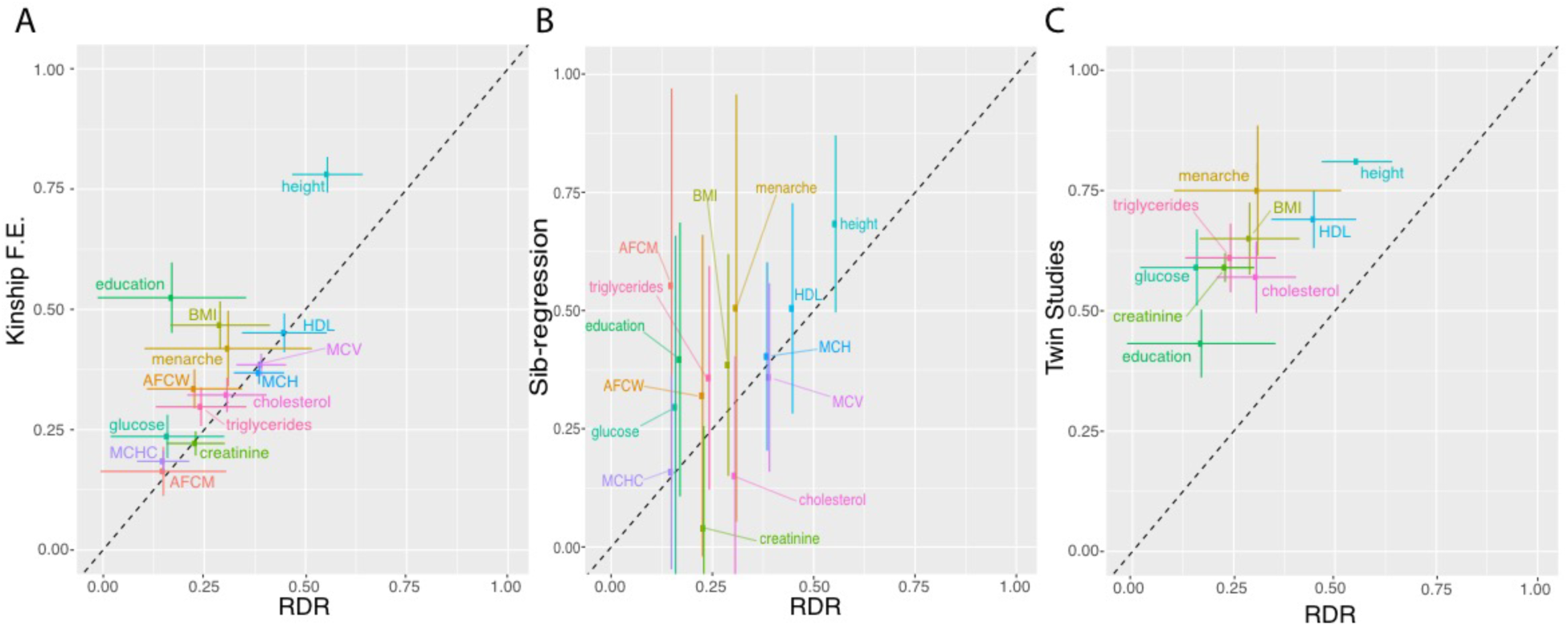
**RDR variance component estimates.** Comparison of RDR heritability estimates with other methods (Table 2). A) Comparison to the ‘Kinship F.E.’ method. Intervals showing +/-1.96 standard errors for the RDR estimates and the ‘Kinship F.E.’ estimates are indicated with the horizontal and vertical bars, respectively. B) Comparison to Sib-Regression^9^ estimates. Intervals showing +/-1.96 standard errors for the Sib-Regression estimates are indicated with the vertical bars. C) Comparison to published twin studies estimates from the Swedish Twin Registry^21^, apart from for education, which is from a meta-analysis of Scandinavian twin studies^22^ (Supplementary Table 5). Trait abbreviations: BMI, body mass index; AFCW, age at first child in women; AFCM, age at first child in men; education, educational attainment (years); cholesterol, total cholesterol; HDL, high density lipoprotein; glucose, fasting glucose; MCH, mean cell haemoglobin; MCHC, mean cell heamoglobin concentration; MCV, mean cell volume.

**Table 2.** **Heritability estimates.** For each trait, the sample size used for the RDR and Kinship methods is given under ‘n’, and the sample size for Sib-Regression (‘Sib-Reg.’) given under ‘sib-pairs’. Each heritability estimate is expressed as a percentage of the phenotypic variance and is followed by its standard error in brackets. RDR and ‘Kinship F.E.’ estimates are from the exact same Icelandic samples with both parents genotyped, whereas Sib-Reg. (Sib-Regression) estimates are from all genotyped Icelandic sibling pairs available. Twin studies estimates are from the Swedish Twin Registry^21^, apart from for education, which is from a meta-analysis of Scandinavian twin studies^22^ (Supplementary Table 5). For the RDR and Kinship F.E. estimates, samples were restricted to those born between 1951 and 1997 for BMI and traits measured from blood, and samples were restricted to those born between 1951 and 1995 for height; Sib-Regression estimates did not have these year of birth restrictions applied. Trait abbreviations: BMI, body mass index; AFCW, age at first child in women; AFCM, age at first child in men; menarche, age at menarche (years); education, educational attainment (years); total chol., total cholesterol; HDL, high density lipoprotein; glucose, fasting glucose; MCH, mean cell haemoglobin; MCHC, mean cell heamoglobin concentration; MCV, mean cell volume.

| Trait | n | sib-pairs | RDR | Kinship F.E. | Sib-Reg. | Twin |
| --- | --- | --- | --- | --- | --- | --- |
| BMI | 19,589 | 56,461 | 28.9 (6.3) | 46.7 (2.5) | 38.5 (12.0) | 65 (3.8) |
| height | 21,802 | 64,847 | 55.4 (4.4) | 78.0 (1.9) | 68.4 (9.6) | 81 (-) |
| AFCW | 22,367 | 30,582 | 22.6 (6.0) | 33.5 (2.1) | 32.0 (17.4) | - |
| AFCM | 17,117 | 21,729 | 14.9 (7.9) | 16.3 (2.6) | 55.3 (21.3) | - |
| menarche | 11,242 | 16,621 | 30.9 (10.5) | 41.9 (4) | 50.6 (23.1) | 75 (6.9) |
| education | 12,035 | 32,542 | 17.0 (9.4) | 52.4 (3.7) | 39.7 (14.8) | 43 (3.6) |
| total chol. | 27,320 | 74,271 | 30.6 (5.0) | 32.2 (1.8) | 15.1 (12.9) | 57 (3.8) |
| HDL | 24,570 | 67,894 | 44.8 (5.3) | 45.1 (2.1) | 50.5 (11.4) | 69 (3.1) |
| triglycerides | 24,099 | 62,746 | 24.2 (5.7) | 29.8 (2.0) | 35.8 (12.1) | 61 (3.7) |
| glucose | 19,500 | 36,469 | 15.9 (7.2) | 23.6 (2.3) | 29.6 (18.5) | 59 (4.0) |
| creatinine | 38,929 | 98,385 | 22.9 (3.7) | 22.2 (1.3) | 4.0 (11.1) | 59 (1.5) |
| MCH | 43,917 | 107,711 | 38.5 (3.2) | 36.8 (1.2) | 40.3 (10.2) | - |
| MCHC | 43,963 | 107,833 | 14.9 (3.3) | 18.4 (1.1) | 15.8 (10.5) | - |
| MCV | 43,919 | 107,702 | 39.1 (3.1) | 38.5 (1.2) | 35.9 (10.2) | - |

For the exact same probands that RDR was applied to, heritability estimates were obtained from the Kinship and Kinship F.E. methods (Methods, Table 2, Figure 2). The Kinship F.E. heritability estimates were 3.9% lower on average than the Kinship heritability estimates, indicating some confounding due to shared family environment in the Kinship estimates. We therefore compared RDR heritability estimates to heritability estimates from the Kinship F.E. method, which we denote as 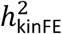. For 11 of the 14 traits, 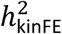 is bigger than 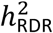 (average 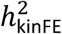 — 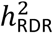 = 12.1%). The traits with the largest difference are educational attainment, height, and body mass index (BMI), with 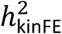 — 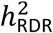 equal to 35.4%, 22.6% and 17.8%, respectively. These results suggest that, for some traits, fitting a family environment effect does not eliminate environmental confounding from Kinship-type methods.

Using Icelandic data, but without limiting to probands with parents genotyped, Sib-Regression estimates of heritability, denoted by 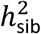, were computed (Methods, Table 2 and Figure 2). Despite having over 100,000 sib-pairs available for some traits, RDR estimates were more precise than Sib-Regression estimates for every trait, and, on average, the standard errors for 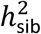 were 2.5 times larger than those for 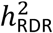.

If a difference between RDR and Sib-Regression exists, it could be a consequence of indirect genetic effects between siblings, or it could be due to epistasis or rare variants. The average of 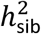 — 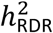 for the fourteen traits is 7.9%, although the difference does not reach significance (*P*=0.071, paired t-test assuming traits independent). For educational attainment, where there is evidence for indirect genetic effects between siblings^15^, 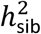 — 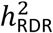 = 22.7%, although the difference does not reach significance (P=0.097, one-sided test assuming estimates are independent).

There are not enough monozygotic twins in the Icelandic data to obtain precise twin estimates of heritability. To compare RDR results with twin studies from a similar population, we took estimates from the Swedish Twin Registry^21^ denoted by 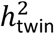, which were available for nine of the fourteen traits (Methods, Table 2, Figure 2, and Supplementary Table 5). The difference 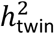 — 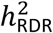 was above zero and statistically significant for all nine traits, with an average difference of 33.2%.

### GREML heritability estimates are biased by indirect genetic effects from relatives

Due to the small population of Iceland, it is not possible to construct the large samples of distantly related people typically used in GREML analyses. To investigate the effect of indirect genetic effects from relatives on GREML analysis, we simulated a population of 20,000 unrelated individuals and their parents (Methods). We simulated a trait determined only by additive, direct effects of SNPs and random noise, and we confirmed that GREML analysis inferred the true heritability, 20%, accurately: mean estimate 20.26% (0.28% S.E.).

Alleles transmitted to offspring are also present in the parents, so have both direct and parental genetic nurturing effects. Let *δ* be the direct effect of a SNP, and let *η* be the parental genetic nurturing effect. The effect of the transmitted allele is therefore (*δ* + *η*). GREML uses only transmitted alleles, so is unable to separate the variance from the direct effect alone, proportional to *δ*^2^, from the variance explained by the combined direct and parental genetic nurturing effects, proportional to (*δ* + *η*)^2^. We investigated this theoretically (Supplementary Text) and by simulating a trait with both direct and genetic nurturing effects. We set the genetic nurturing effect of each variant to be one third of its direct effect, similar to the estimated ratio for educational attainment in Iceland^15^. The direct effects explained 20% of the phenotypic variance. The variance explained by genetic nurturing effects of transmitted alleles alone was 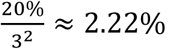. However, the total variance explained by the alleles transmitted to the offspring is much larger than 20%+2.22%=22.22%. This is because the variance explained by a transmitted allele is proportional to 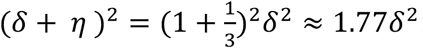. Even though the genetic nurturing effect is only one third of the direct effect, it has magnified the variance explained by the transmitted allele to 1.77 times the variance explained by its direct effect alone. The total variance explained by transmitted alleles is 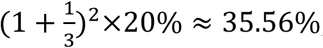, much larger than the heritability, 20%.

The mean heritability estimate from GREML analysis in GCTA was 35.68% (0.27% S.E.). In this simple scenario, GREML analysis estimates the total variance explained by the combined direct and parental genetic nurturing effects of transmitted alleles, rather than the heritability. In contrast, RDR estimates of heritability were approximately unbiased for the simulated traits with genetic nurturing effects (Table 1 and Supplementary Table 3).

## Discussion

We introduced RDR, a novel heritability estimation method, and used it to estimate heritability for 14 quantitative traits in Iceland. Through mathematical investigations and simulations, we demonstrated that RDR estimates of heritability have negligible bias due to environment. RDR estimates may omit some heritability due to rare variants, although the omission is likely to be small unless very rare variants (<0.1% frequency) explain a substantial fraction of the heritability.

Our simulations showed that, in certain scenarios, GREML methods could be used to estimate the variance explained by the combined direct and genetic nurturing effects of transmitted alleles, rather than the heritability. There is evidence for a substantial contribution of parental genetic nurturing effects to educational attainment^15^, implying GREML estimates of the heritability of educational attainment^23^ include variance from parental genetic nurturing effects. In contrast, the RDR estimate of the heritability of educational attainment (17.0%, S.E. 9.4%) does not include variance from parental genetic nurturing effects.

The RDR method requires parents of probands to be genotyped. Large datasets with this property are currently rare, which is the main reason our current study is limited to the Icelandic population. While we cannot rule out the possibility that the heritability of a trait in Iceland is substantially different from the heritability in other European populations, the fact that the effects of many genetic loci are similar between Iceland and other European populations^24-26^ argues against there being major differences in genetic architecture. Nevertheless, some of the difference between 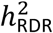 and 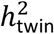 could be due to differences in heritability between our Icelandic sample and the Swedish twin samples. Additionally, some of the difference could be due to overestimation of heritability by twin studies, or due to very rare variants (<0.1% frequency) explaining a substantial fraction of the phenotypic variance. Presently, our data do not allow us to determine the relative contribution of the different possible explanations.

Importantly, our results show that having parental genotypes is a way to obtain precise heritability estimates with negligible environmental bias. In the future, as large cohorts with parents and offspring genotyped become more common, we expect RDR and related methods to become more widely applied. In the more distant future, extensions could include grandparents and other relatives, enabling improved estimation of genetic nurturing effects^15^. This will give a better understanding of how genetic variation affects the social environment across generations, which could change our perspective on the nature-versus-nurture debate.

## Acknowledgements

A.Y. was supported by a Wellcome Trust Doctoral Studentship (099670/Z/12/Z) for part of this project.

## Methods

### Icelandic Sample

All participating subjects donating biological samples signed informed consents and the study was approved by the Data Protection Commission of Iceland (DPC) and the National Bioethics Committee of Iceland. Personal identities of the phenotypes and biological samples were encrypted by a third party system provided by the Icelandic Data Protection Authority.

The Icelandic samples were genotyped using Illumina microarrays as previously described^27^. The whole genomes of 2,636 Icelanders were sequenced using Illumina technology to a mean depth of at least 10X (median 20X)^27^. A total of 35.5 million autosomal SNPs and indels were identified using the Genome Analysis Toolkit version 2.3.9.

The deCODE genealogy database is a comprehensive database that includes information on more than 800,000 Icelandic individuals, deceased and living, dating back to the settlement of Iceland 1,200 years ago. The database is constructed from a nationwide census, conducted regularly from the year 1700, church books and other available information, and is particularly complete for the last 200 years. The database includes, when known, information on parents of each individual, gender, year of birth and, if applicable, year of death.

We restricted our analyses to genotyped individuals with both genetic parents genotyped and all four grandparents in the deCODE genealogy database. This left 54,888 individuals.

The individuals and their parents had all been phased and segments shared identical-by-descent (IBD), both within and between individuals, determined by long-range phasing^28,29^. To reduce bias due to segments incorrectly called as identical-by-descent, we restricted our analyses to segments of length greater than 5 centi-Morgans.

As a measure of ascertainment bias, we compared years of education between the individuals with both parents genotyped and the full set of individuals with education data. Mean years of education for those with both parents genotyped was 15.07 compared to 13.63 for the whole sample with education data. Part of this is due to the fact that those with both parents genotyped were born later than average, and mean levels of education have increased over time. After regressing out year-of-birth (YOB), YOB^2^, YOB^3^, the sample with both parents genotyped still had 0.32 years more education on average, compared to a standard deviation of 3.39 years. This shows that our results are slightly biased towards those with higher socioeconomic status, which, for many traits, is expected to increase heritability^30,31^.

### Trait measurements

As a measure of educational attainment, we used information on years of schooling, available for 63,508 individuals, that originated from questionnaires administered in deCODE’s various disease projects and from routine assessments of elderly nursing home residents. As the data have been gathered over the years for the purpose of descriptive demographics rather than for phenotype use, the questions were originally not standardized across projects and many of them have categorical responses. For this study, to make it as consistent as is possible when it comes to the educational attainment trait studied in the published meta-analysis^32^, efforts were put into mapping the responses to the questionnaires into the UNESCO ISCED classification (http://www.uis.unesco.org/Education/ISCEDMappings/Pages/default.aspx). In particular, the final quantitative measure used, before sex and year-of-birth adjustments, ranges from a minimum of 10 years to a maximum of 20 years.

Height and body mass index (BMI) information, collected primarily through deCODE’s genetic studies on cardiovascular disease, obesity and cancer, were available for 89,615 and 77,285 adult individuals, respectively^25,26^. About 20% of the information was self-reported.

Blood measurements were collected from three of the largest laboratories in Iceland: Landspítali - The National University Hospital of Iceland, Reykjavík; The Laboratory in Mjódd, Reykjavík; Akureyri Hospital, The Regional Hospital in North Iceland, Akureyri; in addition to the Icelandic Heart Association. For many individuals, multiple blood samples had been taken at different time points. To aid comparability with other studies that have used one time-point only, we took only the first measurement of each individual.

Information on ‘age at first child’ (AAFC) was extracted from the deCODE genealogy database. Age at menarche was determined by the answer to the question ‘How old were you when your menstruation started?’ as detailed elsewhere^33^.

Apart from educational attainment, traits were quantile-normalised within each sex. Educational attainment was not quantile-normalised as the measurements fall into discrete categories of years of education. The traits were regressed on year-of-birth (YOB), sex, YOB, YOB^2^, YOB^3^, and the interactions of sex with YOB, YOB^2^, YOB^3^. The residuals of this regression were then used as the phenotype, Y, when fitting the models described below. To ensure our heritability estimates correspond to the adult but not elderly population, we further restricted our analysis to those born between 1951 and 1995 for height, and between 1951 and 1997 for BMI and the traits measured from blood. (Note that for Sib-Regression, the year of birth restrictions were not applied to maximise the sample size.)

### Identification of siblings

For the Sib-Regression estimator, we obtained the relatedness for all pairs of genotyped individuals who share both parents in the genealogy. To ensure we only used true full-siblings, we clustered the pairs by relatedness into four clusters using k-means clustering: unrelated, half-sibling, full-sibling, and monozygotic twins. This left 127,264 full-sibling pairs, comprised of 70,317 unique individuals, whose relatedness distribution had a mean of 0.502 and a standard deviation of 0.0382. To maximise the precision of the Sib-Regression estimator, we did not restrict by age or by number of parents genotyped, so the sample used is different to the sample used for the other estimators.

### Simulation of GREML inference

To simulate SNP based GREML inference on unrelated individuals, we first simulated a population of 20,000 unrelated parent pairs, each comprised of a mother and a father. To do this, we simulated genotypes at 10,000 independent SNPs for each parent, where the maternal/paternal genotype for each SNP was drawn from a Bernoulli(0.5) distribution, giving genotypes for each parent as either 0/0, 0/1, 1/0, or 1/1. For each parent pair, we simulated one offspring by simulating Mendelian transmission independently at each SNP. This gave a population of 20,000 offspring from independent and unrelated parent-pairs.

To compute the SNP based kinship matrix, we first standardised each SNP to have mean zero and variance 1. Let *G* represent the standardised genotypes of the offspring. We calculated the SNP kinship matrix as *R*_snp_ = 10^−4^*GG*^*T*^.

We simulated 10 independent traits determined by additive genetic effects and noise. For each trait, we simulated a normally distributed vector of effects for the 10,000 SNPs: *β∼N*(0, *I*). The additive genetic component of the trait, *A*, was then calculated as *A = Gβ*, scaled so that A had sample mean 0 and sample variance 1. The noise component was simulated as *∊∼N*(0, *I*). The ‘additive’ trait was simulated as 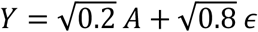.

We simulated 10 independent traits with both additive and genetic nurturing effects. In addition to an additive genetic component simulated as above, we also simulated a genetic nurture component. Let *G*_par_ represent the matrix of standardised parental genotypes, where the parental genotype is defined as the sum of mother’s 0,1,2 genotype and the father’s 0,1,2 genotype. The genetic nurturing component of the trait, *A*_par_, was then calculated as *A*_par_ = *G*_par_β, where *β* is the same vector of effects as for the direct, additive component, *A = Gβ*. We scaled *A*_par_ so that it had sample mean 0 and sample variance 1. The noise component was simulated as *∊∼N*(0, *I*). The trait with genetic nurturing effects was then simulated as

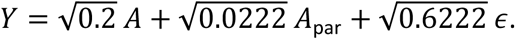

GREML analysis performed on the traits using restricted maximum likelihood in *GCTA*^34^ with *R*_snp_ as the only genetic relatedness matrix.

### Simulations using deCODE data

The deCODE sample has been genotyped on different genotyping arrays, and genotypes have been imputed for millions of variants^27^. To enable us to have a shared set of genotypes for both parents and offspring for a sample of 10,000 individuals with both parents genotyped, we used the imputed genotype data. For all traits other than the ‘rare SNPs’ trait, we used imputed genotypes at the ∼600,000 SNPs that comprise the Illumina Framework SNP set, which is a set of SNPs shared between many of the Illumina genotyping arrays used to genotype the Icelandic sample^27^. We chose these SNPs because they have been imputed with high accuracy: the median imputation information was 0.9999. We further filtered the SNPs so that the minimum imputation information was 0.9999, removing around half of the SNPs. Out of the remaining SNPs passing the filter, we randomly sampled 10,000 SNPs to use as the causal SNPs in our simulations. In the 10,000 selected SNPs, the median imputation information was 1.0000, the minimum minor allele frequency (MAF) was 0.52%, and the median MAF was 22.8%. For the ‘rare SNPs’ trait, we randomly sampled SNPs from all imputed SNPs with MAF between 1% and 0.1% and with imputation information at least 0.9999 and p-value for Hardy-Weinberg deviation greater than 0.05. We sampled 100 such SNPs from each chromosome, giving 2,200 SNPs in total.

For each type of trait, we simulated 500 independent replicates. Each trait had a direct, additive genetic component that explained 40% of the phenotypic variance, which we describe the simulation of here. Apart from for the ‘rare SNPs’ trait, we standardised genotypes so that each SNP’s genotype vector had sample mean zero and sample variance one. Let *G* represent the matrix of standardised genotypes at the 10,000 causal SNPs. We sampled additive effects of SNPs from a normal distribution. Let *β∼N*(0, *I*) represent the vector of SNP effects. The additive genetic component, *A*, was then calculated as *A = Gβ*. The noise component was simulated as *∊∼N*( 0, *I*). The additive genetic component was scaled to have sample variance 1. The ‘additive’ phenotype was then simulated as: 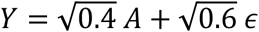. The same process was used for the ‘rare SNPs’ trait, except genotypes were not standardised because the standardisation becomes highly sensitive to fluctuations in estimated allele frequencies for rare SNPs.

For the ‘epistatic’ trait, we simulated a genetic component due to pairwise interactions between SNPs. To do this, we sampled 100 SNPs from the 10,000 SNPs given additive effects. We formed pairwise interaction variables for all pairs of SNPs from the 100 selected SNPs by multiplying the standardised genotypes of each pair of SNPs together. Let *G*_epi_ be the resulting matrix of SNP-SNP interaction variables. We standardised the columns of *G*_epi_ so that each column had sample mean zero and sample variance one. We simulated interaction effects from a normal distribution, *β*_epi_∼*N*(0, *I*), and we formed the pairwise interaction genetic component as *A*_epi_ = *G*_epi_*β*_epi_, and standardised *A_epi_* so that it had sample mean zero and sample variance one. The ‘epistatic’ trait was then formed as:

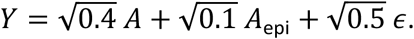

For the ‘regional’ trait, we gave each of the 22 regions of Iceland (called ‘syslas’) a different normally distributed effect, and we scaled the overall variance explained by variation in sysla to be 20% of the phenotypic variance. The phenotype was thus simulated as: 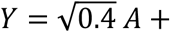 sysla + 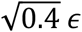, where sysla represents the vector of sysla effects.

For the ‘maternal environment’ trait, we added an environmental effect that was shared between individuals who shared mothers according to the deCODE genealogy database. The effect due to each mother was drawn from a normal distribution, and resulting vector of effects due to maternal environment, M, was scaled to have variance 0.4. The phenotype was simulated as:

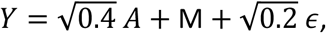

Note, if *Y*_1_ and *Y*_2_ had the same mother in the deCODE genealogy database, then their maternal environment variables were the same.

For the ‘genetic nurturing’ trait, we simulated a component reflecting parental genetic nurturing effects^15^: each genetic variant in the parents was also given an additive effect on the trait of the proband. The additive genetic component was *A = Gβ*, scaled to have variance 1. Let *G*_par_ be the matrix of standardised parental genotypes, where the parental genotype is defined as the sum of the mother’s genotype and the father’s genotype. Then the genetic nurturing component was simulated as *A*_par_ = *G*_par_*β*, scaled to have sample variance 1. This implies that the parental genetic nurturing effects differ only by a constant scale factor from the direct effect of the genetic variant in the offspring. The phenotype was then simulated as

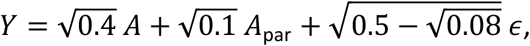

where the scale factor for the residual variance was calculated so that the total phenotypic variance, which includes the covariance between *A* and *A*_par_, was 1. For this trait, *v*_*e∼g*_ = 0.1 and 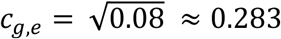.

### Selection of estimates from twin studies

There are not enough monozygotic twins in Iceland to give precise twin based estimates of heritability, forcing us to compare our results with external twin studies. The Swedish Twin Registry^21^ is a large sample of twins from a population of similar cultural and genetic composition to Iceland, giving the most precise and valid comparison possible based on published data^35-39^. The exception is for education, where we used a meta-analysis of Scandinavian twin studies for increased precision^22^. For BMI and traits measured from blood, unlike our estimates, the Swedish Twin Registry estimates did not exclude elderly individuals. This is unlikely to account for the higher estimates in the Swedish Twin Registry, as twin correlations and heritability estimates are generally lower in the elderly population^2^.

We took the heritability estimate from the additive-common-environment (ACE) model^2,3^ when provided. ACE estimates were not provided for the blood lipid traits, but monozygotic and dizygotic twin correlations were^38^. We used these to obtain the moment based estimate of the heritability under the ACE model by the formula: 2(*r*_MZ_ – *r*_DZ_), where *r*_MZ_ is the phenotypic correlation for monozygotic twins, and *r*_DZ_ is the phenotypic correlation for dizygotic twins. We took the weighted average of the same-sex and opposite-sex dizygotic twin correlations to estimate *r*_DZ_. For creatinine, the ACE estimate was not provided, and neither were the twin correlations, so we took the published heritability estimate from the ADE model (additive-dominance-environment). The studies used and methods used are summarised in Supplementary Table 5. For height, heritability estimates were only provided for males and females separately, so we took the average estimate. The standard error was not provided. Height and weight estimates were based on self-reported data, whereas our estimates were based on approximately 80% measured and 20% self-reported data. This would be expected to increase our heritability estimates for height and BMI relative to the twin estimates due to a reduction in measurement error. For education, we used a meta-analysis of twin studies in Scandinavian countries, including Sweden, to give a more precise estimate^22^. We could not find published estimate based on the Swedish Twin Registry for the haemoglobin traits and for age at first child, so we excluded them from the comparison.

### Calculation of relatedness matrices

To calculate *R, R*_par_, and, *R*_o,par_, we used formulae based on the genetic covariance in a population descending from a finite number of ancestors^40^ (Supplementary Note):

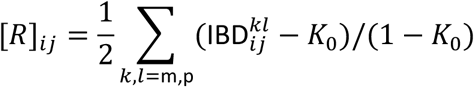

where *K*_0_ is the mean kinship over all pairs in the population, and 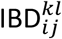 is the proportion of the maternally inherited haplotype of *i* shared identical-by-descent (IBD) with the paternally inherited haplotype of *j*;

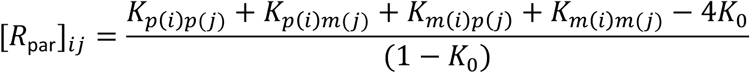

where *K*_*p*(*i*)*m*(*j*)_ is the kinship between the father of *i* and the mother of *j*;

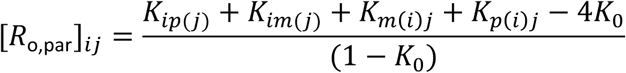

where *K*_*im*(*j*)_ is the kinship between *i* and the mother of *j*, etc. The statistics of these matrices are recorded in Supplementary Tables 6 and 7.

### Model Fitting

To obtain heritability estimates from the Kinship method, we fit the following model for a vector of phenotype observations ***Y***:

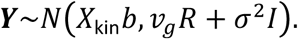

For the Kinship F.E. model, we added a variance component that modelled shared family environment:

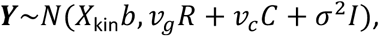

where [*C*]_*ij*_ = 1 if *i* and *j* shared a mother according to the deCODE genealogy database, otherwise [*C*]_*ij*_ = 0. For all of the simulated traits other than the ‘regional’ trait, *X*_kin_ was a constant. For the ‘regional’ trait, it also included the top 20 genetic principal components. In the real trait analysis, *X*_kin_ included the top 20 genetic principal components. For both the Kinship and Kinship F.E. methods, we estimated model parameters by unconstrained restricted maximum likelihood in *GCTA*^34^.

The relatedness disequilibrium regression (RDR) covariance model is

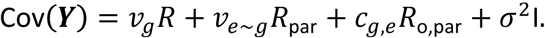

We investigated fitting this model by least squares regression of the off-diagonal elements of the sample phenotypic covariance matrix on the off-diagonal elements of the relatedness matrices:

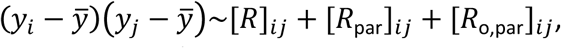

where *y*_*i*_ is the phenotype observation for individual *i*, and 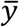 is the sample phenotype mean. We excluded both parent-offspring and grandparent-grandchild pairs from the regression, as these pairs violate the relationship between [*R*]_*ij*_ and [*R*_par_]_*ij*_ required for removal of environmental bias from estimation of *v*_*g*_ (Figure 1 and Supplementary Note). These pairs comprised around 0.0014% of all the pairs we could have used. We also investigated fitting the model by unconstrained restricted maximum likelihood in *GCTA*^34^, under the assumption the trait follows a multivariate normal distribution:

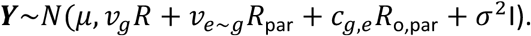

For the maximum likelihood method, one can only remove individuals, and all the pairs including that individual, not arbitrary pairs. Around 30% of the sample with both parents genotyped have an ancestor who also has both parents genotyped. We therefore did not exclude individuals so that no parent-offspring and no grandparent-grandchild pairs were present, as this would have resulted in a large loss of sample size.

In our simulations, we found that RDR estimates from maximum likelihood and RDR estimates from least-squares were both approximately unbiased, with no consistent advantage in bias evident from fitting the model by least-squares after excluding parent-offspring and grandparent-grandchild pairs (Supplementary Table 1). However, least-squares estimates were considerably less precise than those from maximum likelihood. We therefore used maximum likelihood without exclusion of parent-offspring and grandparent-grandchild pairs (which comprised only 0.0014% of the pairs with both parents genotyped) for all analyses in the main text.

To obtain Sib-Regression estimates^9^, we fit the regression model

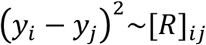

for all *i,j* such that *i* and *j* are full-siblings. We fit the regression model by least-squares using custom *R* code. The estimate of *v*_*g*_ is then minus one half of the estimated regression coefficient. We compared estimating standard errors by the approximate formula given in the original Sib-Regression paper^9^ (equation 17) and estimating standard errors by treating Sib-Regression as a standard univariate linear regression with uncorrelated observations. For the ‘additive’ simulated trait, both gave almost exactly the same estimated standard error, which underestimated the standard error by approximately 9%. We used standard errors estimated from treating sib-regression as a standard univariate linear regression with uncorrelated observations for all other results.

